# An Optimization framework for Nonspiking Neuronal Networks and Aided Discovery of a Model for the Elementary Motion Detector

**DOI:** 10.1101/666149

**Authors:** Arunava Banerjee

**Affiliations:** Computer and Information Science and Engineering, University of Florida, PO Box 116120, Gainesville, FL 32611, USA

## Abstract

We present a general optimization procedure that given a parameterized network of nonspiking conductance based compartmentally modeled neurons, tunes the parameters to elicit a desired network behavior. Armed with this tool, we address the elementary motion detector problem. Central to established theoretical models, the Hassenstein-Reichardt and Barlow-Levick detectors, are delay lines whose outputs from spatially separated locations are prescribed to be nonlinearly integrated with the direct outputs to engender direction selectivity. The neural implementation of the delays—which are substantial as stipulated by interomatidial angles—has remained elusive although there is consensus regarding the neurons that constitute the network. Assisted by the optimization procedure, we identify parameter settings consistent with the connectivity architecture and physiology of the Drosophila optic lobe, that demonstrates that the requisite delay and the concomitant direction selectivity can emerge from the nonlinear dynamics of small recurrent networks of neurons with simple tonically active synapses. Additionally, although the temporally extended responses of the neurons permit simple synaptic integration of their signals to be sufficient to induce direction selectivity, both preferred direction enhancement and null direction suppression is necessary to abridge the overall response. Finally, the characteristics of the response to drifting sinusoidal gratings are readily explained by the charging-up of the recurrent networks and their low-pass nature.

## Introduction

A major goal of Neuroscience is to understand how the activity of identified neural circuits relate to behavior. Recent advances in (semi)automated reconstruction from EM data and immunolabeling have generated a wealth of information regarding the neural connectivity architecture and polarity of synapses in the brains of model organisms. For example, [33] has released an EM volume of the complete Drosophila brain, parts of which have been reconstructed for neural connectivity. These notable advances have, however, not been followed by the anticipated spate of neural implementation solutions to well characterized high level operations. This discrepancy stems from the fact that the dynamics of a neural circuit is determined not only by connectivity and synaptic polarities, but also by the synaptic gain profiles, information which the current techniques do not reveal at the necessary level of detail. Lacking synaptic profile information, relating circuits to behavior has therefore been difficult.

The observation that network behavior is influenced by synaptic profiles also implies that it is, in principle, possible to infer synaptic profiles from the recorded behavior of a network. Such a data driven approach, where a parameterized model is adapted to the characteristics of a dataset, has been the cornerstone of recent successes in Machine Learning. The general problem in the current context, where a network comprises both spiking and nonspiking neurons, is likely both formally and computationally intractable. However, as we demonstrate in this article, a more restricted class, that of nonspiking neuronal networks, lends itself to this approach. Specifically, we show that local analysis of the trajectories of the corresponding dynamical system is tractable: in operational terms, given a parameterized network of compartmentally modeled conductance based nonspiking neurons interacting with tonically active synapses, computational optimization can tune the parameters to cause the network’s behavior to display desired properties. The optimization procedure developed is general and applies to a large class of parameters, including synaptic profiles as well as morphological properties of the constituent neurons.

Nonspiking neuronal networks abound in the sensory and motor periphery, the set of circuits composed of photoreceptors, horizontal, bipolar, and amacrine cells in the vertebrate retina [10] being among the most widely cited examples. In order to showcase the strength of the optimization driven approach, we have however chosen a network whose (a) architecture is conserved across multiple insect species, (b) has been characterized in substantial detail, and (c) has well established theoretical models whose relationship to the neural implementation remains to be reconciled. In resolving this problem, we show that computational optimization of parameterized synaptic profiles to fit a network’s behavior can be a potent tool when combined with connectivity and synaptic polarity information.

The elementary motion detector (EMD) in the fly brain is a paradigmatic neural computation that has been the subject of intense investigation over decades [6]. Established theoretical models, the Hassenstein-Reichardt (HR) [15] and the Barlow-Levick (BL) [4] detectors prescribe delay lines, the neural implementation of which has thus far remained elusive. Much is however known about the neural circuits that implement the EMD, that we summarized here.

The compound eyes of flies consist of anatomically identical units, called omatidia, laid out in a hexagonal lattice. Visual information processing begins at the photoreceptors in the omatidium, advancing thereafter through neurons in four retinotopically organized neuropile, the lamina, medulla, lobula, and lobula plate. Broad interest in fly motion vision, stemming from its status as a canonical computation, has led to the accumulation of a wealth of data— particularly with regard to the Drosophila—concerning the connectivity architecture and physiology of the neurons in the repeating modules associated with each omatidium, in the lamina [20, 30] as well as the medulla and lobula [29, 26]. Briefly, axons of photoreceptors R1-R6 innervate lamina monopolar cells (LMC) L1-L3 in the corresponding lamina module. Of the cells identified in the lamina, only the joint silencing of the similarly responding [8] L1 and L2 abolishes direction selectivity [30]. Exiting the lamina, motion information is extracted in parallel pathways with L1 feeding the brightness increment (ON) and L2 feeding the brightness decrement (OFF) circuitry [16]. The connectome of the ON module in the medulla has been elucidated in substantial detail [29]. Based on the preponderance of different synaptic contacts, the following core circuit emerges (Fig.1a). The primary targets of L1 are Mi1, Tm3, L5, and C3. L5 drives Mi4, C3 drives Mi9, and Mi4 and Mi9 are reciprocally connected. Lastly, Mi1, Tm3, Mi4, and Mi9 constitute the primary inputs to T4 which is the first cell on the pathway to exhibit direction selectivity. Although Fig.6 in [29] contains additional cells and synaptic contacts, our results demonstrate that this parsimonious circuit is sufficient to manifest direction selectivity, with Mi1/Tm3 providing the direct input and Mi4/Mi9 providing the delayed input into T4. The narrow receptive field T4 comes in four flavors: T4a-d each tuned to one of four cardinal directions. Whole-cell recordings [5] indicate that all of the noted cells are graded potential neurons, and therefore communicate using tonically active synapses [17]. The connectome of the OFF module exhibits strong parallels [26].

**Figure 1:**
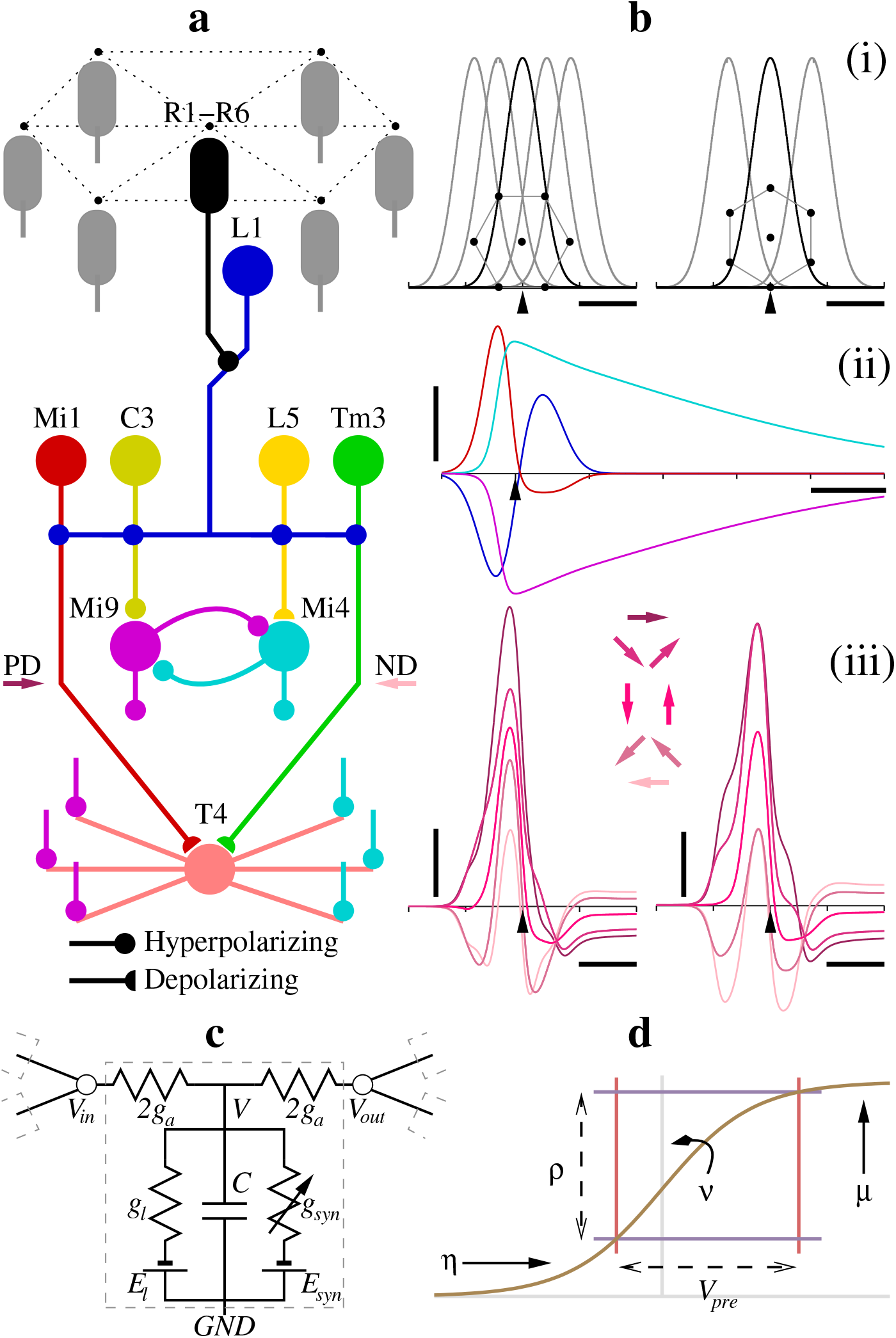
Neural circuit and representative behavior. **a,** Circuit for a single module. The circuit is replicated for all modules in the hexagonal lattice, each receiving input from its corresponding photoreceptors (gray). The home module T4 receives input from three Mi9 (from contiguous neighboring modules), three Mi4 (from symmetrically opposing modules), and the Mi1 and Tm3 of the home module. Synaptic parameters, set symmetrically for each Mi4/Mi9 opposing pair, differ for T4s tuned to different cardinal directions (Methods). **b,** Responses of neurons to a 2° wide bar of light traveling at 12°/*s*. Responses from circuits tuned to two cardinal directions are shown; responses for the other two cardinal directions are symmetric. (i) Temporal profiles of the normalized stimulus intensity sensed by the photoreceptors of seven contiguous omatidia in the lattice for the two cardinal circuits. Insets display the orientation of the lattice points with respect to the bar stimulus traveling left to right. (ii) Responses of L1, Mi1, Mi4, and Mi9, color coded and shifted along the ordinate by their respective equilibrium resting potentials. The responses of Mi4 and Mi9 are also scaled ×10. (iii) Responses of the home module T4s tuned to the two cardinal directions. Stimulus directions PD, PD±*π*/4, PD±*π/*2, ND±*π/*4, and ND, are with respect to the insets in (i). Horizontal scale bars in (i), (ii), (iii) = 500*ms*. Vertical scale bar in (ii) = 5*mV*, in (iii) = 1*mV*. Pointers on abscissa mark time alignment for the home module. **c,** Conductance based model for a single compartment. Linked compartments *i, j* satisfy 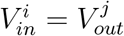. **d**, First model of synapse.

We chose to computationally model the ON pathway because data pertaining to the dimensions of neurites and the polarity of synapses is largely available in this case [29]. The principles we have discovered, however, apply to the OFF pathway as well.

## Results

### Parameterization of Network and Intractability of Global Analysis

We begin with the standard conductance based compartmental modeling methodology, where each neuron is partitioned into equipotential segments whose terminals are then linked to assemble the network (Fig.1c). Each compartment *i* is modeled using the differential equation,

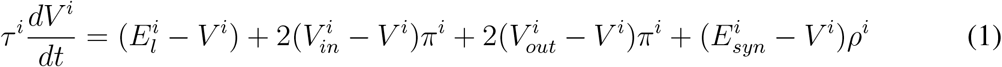

where 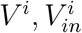 and 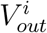 denote the time varying membrane potential at the middle and the two terminals of a compartment, and 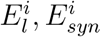 denote the constant leak and constant synaptic potentials. Also, 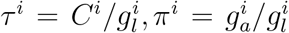, and 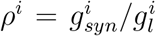, where 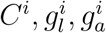 and 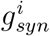 denote the constant capacitance, constant leak, constant axial, and time varying synaptic conductances. The scaling by 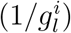 makes *π*^*i*^ and *ρ*^*i*^ dimensionless quantities, which in turn aid in the inter-pretability of the equation.

Compartments are linked as mandated by the modeled cellular morphology of the neurons (Fig.1a and c). Each terminal of each compartment satisfied one current balance algebraic constraint: if the terminal corresponding to 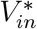 of compartment ∗ is linked to the terminals corresponding to 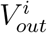 of compartments *i* = 1 … *m*, then setting 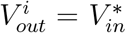 for all *i* = 1 … *m*, we have

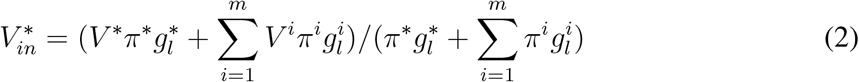

and likewise, if the terminal corresponding to 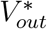 of compartment ∗ is linked to the terminals corresponding to 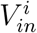 of compartments *i* = 1 … *m*, then setting 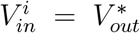 for all *i* = 1 … *m*, we have

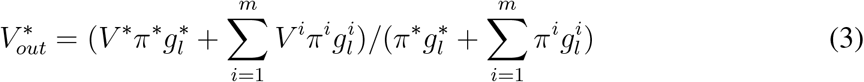

The 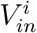 and 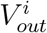 in Eq. 1 can be eliminated by plugging in Eq. 2 and Eq. 3, resulting in a system of ordinary differential equations (ODE) involving only the *V*^*i*^ of each compartment.

It is instructive at this juncture to note that the simplest form this system of equations can take is when the time varying trajectories of the synaptic drives, *ρ*^*i*^, are predetermined independent of the *V*^*i*^s, such as in a feedforward network where the dynamics of the system is solved layer by layer. Even so, what results is a nonhomogeneous linear time-varying system of ODEs of the form **V**′ = **A**(*t*)**V**+ **b**(*t*), where **V**′ is the vector of time derivatives of the *V*^*i*^ of the compartments, **A**(*t*) is a time varying matrix whose elements are composed of *π*^*i*^, 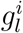, and the time varying *ρ*^*i*^, with the structure of **A**(*t*) determined by the connectivity between the compartments, and **b**(*t*) is a time varying vector with elements composed of 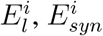 and *ρ*^*i*^.

The general solution to **V**′ = **A**(*t*)**V**+ **b**(*t*) is achieved [23] by solving for the state transition matrix **Φ**(*t, τ*) for the homogeneous system **V**′ = **A**(*t*)**V** using the Peano-Baker series

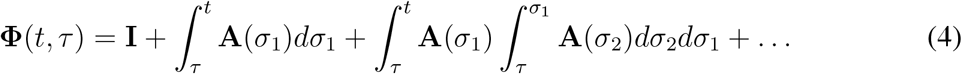

and then incorporating **b**(*t*) to arrive at the solution for initial condition **V**(*t*_0_) given by

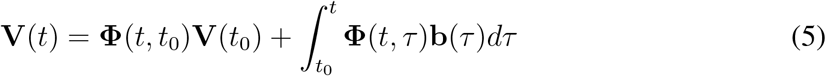

In cases where the fundamental matrix **Ψ**(*t*), satisfying the matrix ODE **Ψ**′(*t*) = **A**(*t*)**Ψ**(*t*) can be computed, **Φ**(*t, τ*) can be solved without resorting to the Peano-Baker infinite series, as **Φ**(*t, τ*) = **Ψ**(*t*)**Ψ**^−1^(*τ*). With the reasonable assumption that **A**(*t*) is periodic with a suitably long period, solving for **Ψ**(*t*) requires the Floquet decomposition of **A**(*t*). Unfortunately, this approach too eludes analytic solutions even for simple periodic parametric functions of time, *ρ*^*i*^, since it involves matrix exponentiation as a function of the parameters [23], among other reasons.

Finally, in the case of recurrent networks, the study of which is our primary objective, the *ρ*^*i*^s can not be predetermined independent of the *V*^*i*^s; each *ρ*^*i*^ is a nonlinear function of a *V*^*j*^, the membrane potential of the appropriate compartment of the presynaptic neuron. The immediate consequence is that the system of ODEs is now nonlinear and none of the above approaches apply. Recognizing then that global analysis of the dynamics of the network is intractable, that is, parameterized closed form solutions may not exist, we are left with the possibility of conducting a local analysis. To elaborate, given a solution trajectory **V**(*t*) of the compartments for some setting of all the parameters, is it possible to characterize solution trajectories in a local neighborhood of those parameter settings? That this is feasible is shown in the next section.

Returning to the network parameterization, what remains is the specification of the model for the time varying relative synaptic conductances 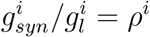 as a function of the presynaptic potential, i.e., *ρ*^*i*^(*V*_*pre*_), where *V*_*pre*_ denotes the *V*^*j*^ of the presynaptic compartment. Noting that the synapses under consideration are tonically active, the time scales at which the synapses operate vis-à-vis the time scale of the network phenomenon under consideration, dictate the choice of the synaptic model. We consider two models. The first is where the network phenomenon operates at a time scale much larger than that of the synapse, so that the synapse can be modeled to be essentially instantaneous. The second is where their time scales are comparable, in which case the synaptic conductance is modeled using an ODE.

In the first model, 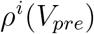 is parameterized as a monotonically increasing and saturating function of the instantaneous potential *V*_*pre*_ of the appropriate presynaptic compartment:

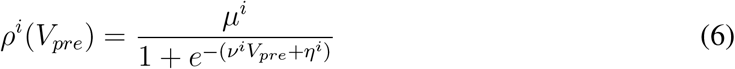

where the parameters *μ*^*i*^, *ν*^*i*^, and *η*^*i*^, set to be strictly positive, determine the gain, sensitivity, and baseline relative conductance (Fig.1d). The second model specifies the time varying synaptic conductance via standard receptor channel kinetics [9]:

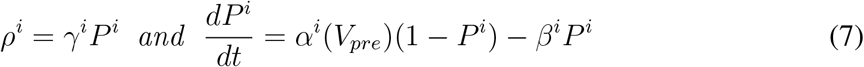

where *γ*^*i*^ denotes the maximum relative conductance, *P*^*i*^ is the time varying fraction of open channels, *β*^*i*^, modeling the closing rate of channels, is a positive constant, and *α*^*i*^, modeling neurotransmitter concentration, is again a monotonically increasing and saturating function of *V*_*pre*_ set as 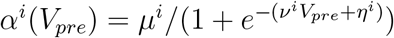.

The entire network is thus specified by the set of parameters 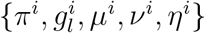 (for the first synaptic model Eq. 6) or 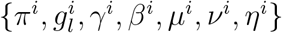 (for the second synaptic model Eq. 7), with *i* ranging over all compartments. We emphasize that the tractability of the upcoming local analysis hinges on each parameter being a scalar quantity so that the entire network is parameterized as a vector in a finite dimensional Euclidean space. In addition, although the analysis applies to all parameters, the morphological parameters, *π*^*i*^ and 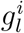, may be fixed based on knowledge of the morphology of the neurons in the circuit under consideration. In the case of the EMD, these were computed for each cylindrical compartment of diameter *d*^*i*^ and length *l*^*i*^ using reported values in [22, 29], and universal membrane constants [12] (Methods). *τ*, being invariant to *d* and *l*, had the value 20*ms* at all compartments. We also set *E*_*l*_ = −55*mV*, and *E*_*syn*_ = −85*mV* for hyperpolarizing and *E*_*syn*_ = 0*mV* for depolarizing synapses.

The input drive into a network is specified as time varying trajectories of presynaptic potentials at the input synapses of the network. Given initial conditions *V*^*i*^(0) (and *P*^*i*^(0) for the second synaptic model) for all compartments *i*, the entire differential-algebraic system of equations traces out a trajectory for *V*^*i*^(*t*), 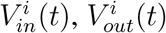 (and *P*^*i*^(*t*)). We restrict initial conditions to respective equilibrium resting potentials (and equilibrium resting fraction of open channels for synapses), thereby eliminating them as free parameters, to focus the analysis solely on the network parameterization. Assuming baseline input drive to be a canonical constant over all time, the network is at its equilibrium resting state when *dV*^*i*^/*dt* = 0 for all compartments and *dP*^*i*^/*dt* = 0 for all synapses. Setting *dV*^*i*^/*dt* = *dP*^*i*^*/dt* = 0 for all *i* turns the differential-algebraic system of equations Eq. 1, Eq. 2, Eq. 3, and Eq. 6 or Eq. 7, into an algebraic system of equations. We initialize this system at 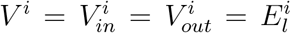 for all *i* (and 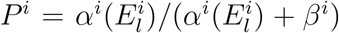 for the synapses) and solve it using a fixed point Jacobi iteration scheme: at each step the *V*^*i*^s are recomputed from the differential-turned-algebraic equations, following which the 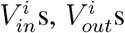, and *P*^*i*^s are recomputed from their respective equations using the new *V*^*i*^s. The iterations are continued until convergence, set as all new values evaluating to within an error bound of their previous values.

We make two final observations. First, although we have used a specific function, the sigmoid, in the synaptic models, the upcoming analysis is general and applies to any class of parameterized smooth functions. Second, voltage gated channels have not been included in the compartmental model ODE to eliminate unnecessary clutter. Mathematically speaking, synaptic channels transform into voltage gated channels when *V*_*pre*_ is replaced by the *V*^*i*^ of its own compartment. The forthcoming analysis is therefore without loss of generalization.

### Local Analysis: Gradient of Voltage Trajectory and Optimization

For any given time varying drive at the input synapses of a network that begins with the canonical baseline value, the differential-algebraic system of equations for the network traces out a corresponding trajectory starting at its equilibrium state. We linearize the system of ODEs with respect to all parameters in a local neighborhood of this trajectory. Assuming infinitesimal perturbations Δ, and dropping higher order terms, we get from Eq. 1, for each compartment *i*, the ODE

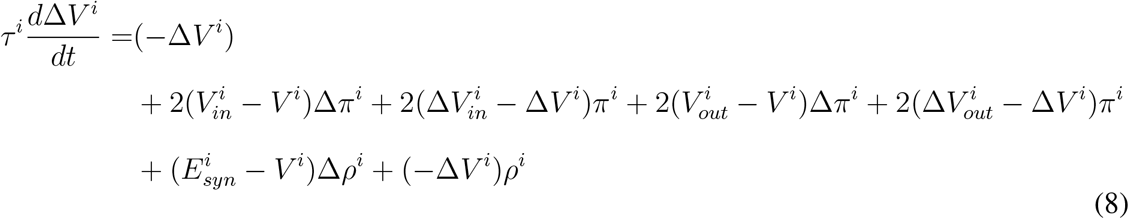

where, for the particular functional form of the first synaptic conductance model Eq. 6,

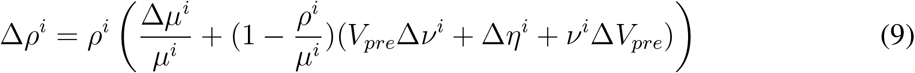

and, for that of the second synaptic conductance model Eq. 7,

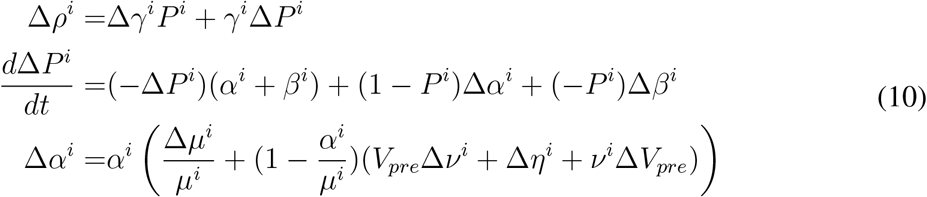

In addition, from the algebraic equations, Eq. 2 and Eq. 3, we get, respectively,

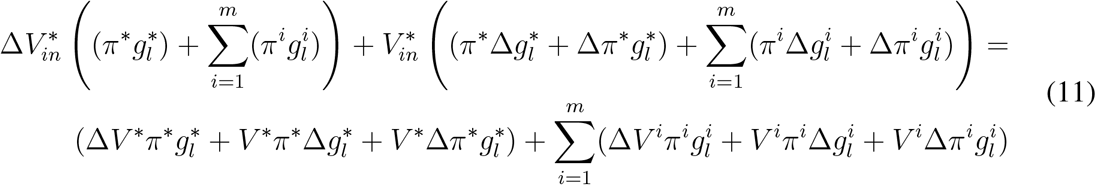

and

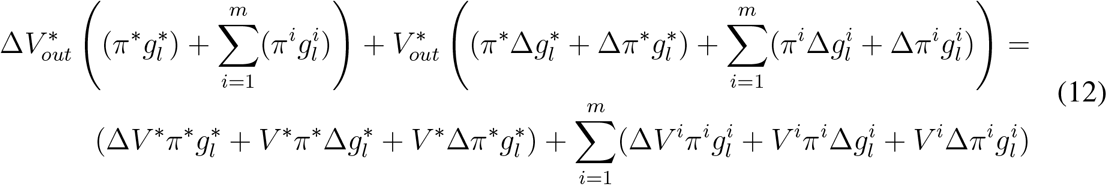

Careful inspection of the system Eq. 8, Eq. 11, Eq. 12, and Eq. 9 (for the first synaptic model) or Eq. 10 (for the second synaptic model) reveals what has been achieved. First, the Δ*V*^*i*^s (and Δ*P*^*i*^s in the case of the second synaptic model) are now defined as a system of linear, albeit nonhomogeneous and time-varying ODEs. Second, the Δ*V*^*i*^s are linear in the parameters, 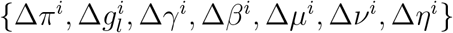.

We can now determine how the trajectory **V**(*t*) composed of the *V*^*i*^(*t*) of all compartments *i*, changes with infinitesimal changes in the parameters that specify a network. Let *X* be a proxy for any of the parameters {π, *g*_*l*_, *γ, β, μ, ν, η*}. Then, 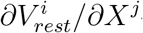—the change in the equilibrium resting potential of *V*^*i*^ per unit change in parameter *X*^*j*^—is obtained by solving the differential-algebraic system of equations with exactly one perturbation Δ*X*^*j*^ = 1 and all other perturbations = 0, setting *d*Δ*V*^*i*^/*dt* = 0 (and *d*Δ*P*^*i*^*/dt* = 0) for all compartments *i*, and setting the input drive to the canonical baseline. That is, 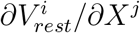 is the fixed point of Δ*V*^*i*^ (calculated as in the previous section) when Δ*X*^*j*^ = 1 is the only nonzero parameter perturbation. Likewise, *δV*^*i*^/*∂X*^*j*^—the time varying change in the trajectory of *V*^*i*^(*t*) per unit change in parameter *X*^*j*^—is obtained by solving the differential-algebraic system of equations for Δ*V*^*i*^, initialized at 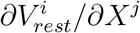, setting that perturbation Δ*X*^*j*^ = 1 and all other perturbations = 0. That is, *δV*^*i*^*/∂X*^*j*^ is the solution to Δ*V*^*i*^(*t*) when initialized at 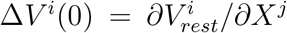 and solved for Δ*X*^*j*^ = 1 as the only nonzero parameter perturbation.

In this general setting, tuning the parameters of a network so as to elicit a desired property of the trajectory **V**(*t*) involves incrementally updating the parameters *X*^*j*^ informed by properties of the trajectory *δV*^*i*^*/∂X*^*j*^. Given an appropriate functional *G* applied to *δV*^*i*^*/∂X*^*j*^, gradient ascent updates can be computed for each parameter *X*^*j*^ as Δ*X*^*j*^ = *ζG*[*δV*^*i*^*/∂X*^*j*^], where *ζ* is the learning rate. The choice of the functional *G*, of course, depends on the specific scenario under consideration. We consider two scenarios; others can be worked out in like fashion.

Let *E*[*f*] be a functional defined on function *f*. Then, the functional derivative, *δE/δf* is, by definition, the linear functional such that *E*[*f* + *ϵg*] − *E*[*f*] = *ϵ* × (*δE/δf*)[*g*] as *ϵ* → 0. This linear functional can be specified in multiple ways: as a function in the integral operator, (*δE/δf*)[*g*] = (*δE/δf*) × *g*, as a point evaluation on a linear map on *g*, etc.

In a scenario where the desired trajectory *U*^*i*^(*t*) of compartment *i* is known for time period [0, *T*], the error functional, *E*[*V*^*i*^], can be defined as:

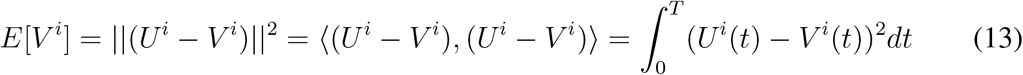

It then follows from the definition above that *δE/δV*^*i*^ is the integral operator specified by −2(*U*^*i*^ − *V*^*i*^), and therefore, 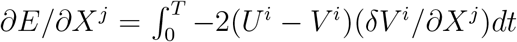. In this case then, the functional *G*[*δV*^*i*^*/∂X*^*j*^] referred to above corresponds to the inner product in function space between the trajectories 2(*U*^*i*^ − *V*^*i*^) and *δV*^*i*^*/∂X*^*j*^, the dropping of the negative sign enforcing descent on *E*.

In the case of the EMD however, the desired trajectory *U*^*i*^(*t*) is unknown. What is known is that the time to maximum (minimum) response of *V*^*i*^(*t*) of the compartments of Mi4 (Mi9) are to be delayed by a specific amount with reference to that of Mi1. Setting *t*\* to denote the desired time to reach the extremum, the error functional *E*[*V*^*i*^] can be defined as:

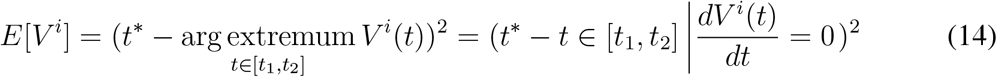

where the second equality follows from the assumption that *V*^*i*^ is unimodal over the range [*t*_1_, *t*_2_] and therefore attains its extremal value at the unique time at which *dV*^*i*^*/dt* = 0.

We can now analyze *E*[*V*^*i*^ + *ϵW*^*i*^] as follows. Let 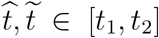 be the times at which 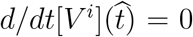 and 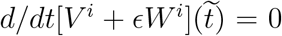. Then applying a Taylor expansion to the second equation around 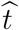 and dropping higher order terms noting that 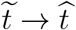 as *ϵ* → 0, we get

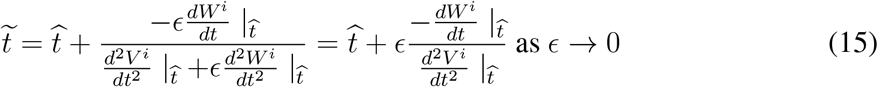

Based on the definition of functional derivative, we then have

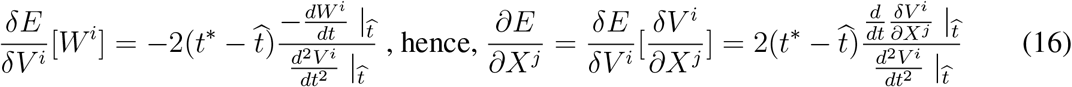

We now make two observations. First, both 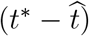 and 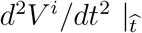 do not depend on the choice of *X*^*j*^ and thus amount to a scaling factor in the gradient. Second, the sign of 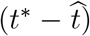 is positive so long as 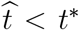, and the sign of 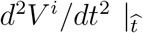 is negative if we choose the argmax of the trajectory of an Mi4 compartment and is positive if we instead choose the argmin of the trajectory of an Mi9 compartment. Therefore, if we choose to push the argmax of the *V*^*i*^ of an Mi4 compartment to the future, the functional 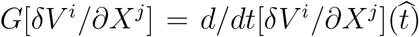, that is, the time derivative of the trajectory *δV*^*i*^*/∂X*^*j*^ at the instant when *V*^*i*^ attains its maximum. The sign is flipped if we choose to push the argmin of the *V*^*i*^ of an Mi9 compartment into the future.

### The Substantial Requisite Delay in the EMD and its Deduced Origin

What makes constructing a model neural circuit implementation of the EMD difficult is not the mere presence of the delay lines as prescribed by the HR and BL models, but the magnitude of the requisite delay as can be inferred from available stimulus-response recordings. The authors in [13], for example, presented a moving 2° ON bar at various orientations and velocities in the receptive field of a T4 cell of a Drosophila, while concurrently recording the T4’s membrane potential response. They found robust directionally selective responses in the behaviorally relevant velocity range of 14° − 112°/*s*. The delay necessary for a robust response at 14°/*s* can be determined from numerical experiments based on anatomical data as described next.

The *∼* 750 omatidia in the Drosophila eye [11] are arranged with interomatidial angle of *≈* 5.1° [27]. The normalized angular sensitivity of photoreceptors R1-R6 in each omatidium is well approximated by 2-D isotropic Gaussians with full width at half maximum (FWHM) estimated at 4.2° when the combined effects of geometric and wave optics are invoked [27]. Fig.1b-i presents numerical simulations of the temporal profiles of the normalized stimulus intensity sensed by the photoreceptors of seven contiguous omatidia in the lattice of the compound eye of the Drosophila for a 2° ON bar traveling at 12°/*s*. The peak to peak delay between the profiles of neighboring photoreceptors for the two lattice orientations are 212 and 368*ms*, respectively, with a FWHM of *≈* 360*ms* for each profile. Clearly, a delay of the order of hundred milliseconds or more would be necessary to induce robust direction selectivity at this stimulus velocity. Such a delay can either arise from complex synaptic dynamics, recurrent network dynamics, or a combination thereof. Our optimized model demonstrates that recurrent network dynamics suffices as a parsimonious explanation (Fig.1b-iii).

Central to the model is a Mi4-Mi9 recurrent network whose synapses, when tuned via the optimization procedure, cause the response of both Mi4 and Mi9 to be delayed and extended in time (Fig.1b-ii). In order to guarantee that the optimization procedure did not simply raise the synaptic time constants to precipitate the delay, we used the first synaptic model Eq. 6 whose response to changes in the presynaptic potential is instantaneous. By eschewing delays, we were assured that the synaptic dynamics did not pose a confound for the delays generated via recurrent network dynamics. Furthermore, the parameters of the network model optimized were restricted to the {*μ*^*i*^, *ν*^*i*^, *η*^*i*^} for all synapses on Mi4 and Mi9; all other parameters were fixed to values determined from the anatomy and physiology of the Drosophila optic lobe. To achieve the delay, we chose the functional *G*[·], introduced in the previous section, to be the time derivative of *δV*^*i*^*/∂X*^*j*^ for a Mi4 compartment at the instant when its *V*^*i*^(*t*) reached its maximum in response to a bar stimulus. This allowed us to make progressive infinitesimal updates to the parameters that pushed Mi4’s and Mi9’s time to peak to the future (Fig.2a-iv).

**Figure 2:**
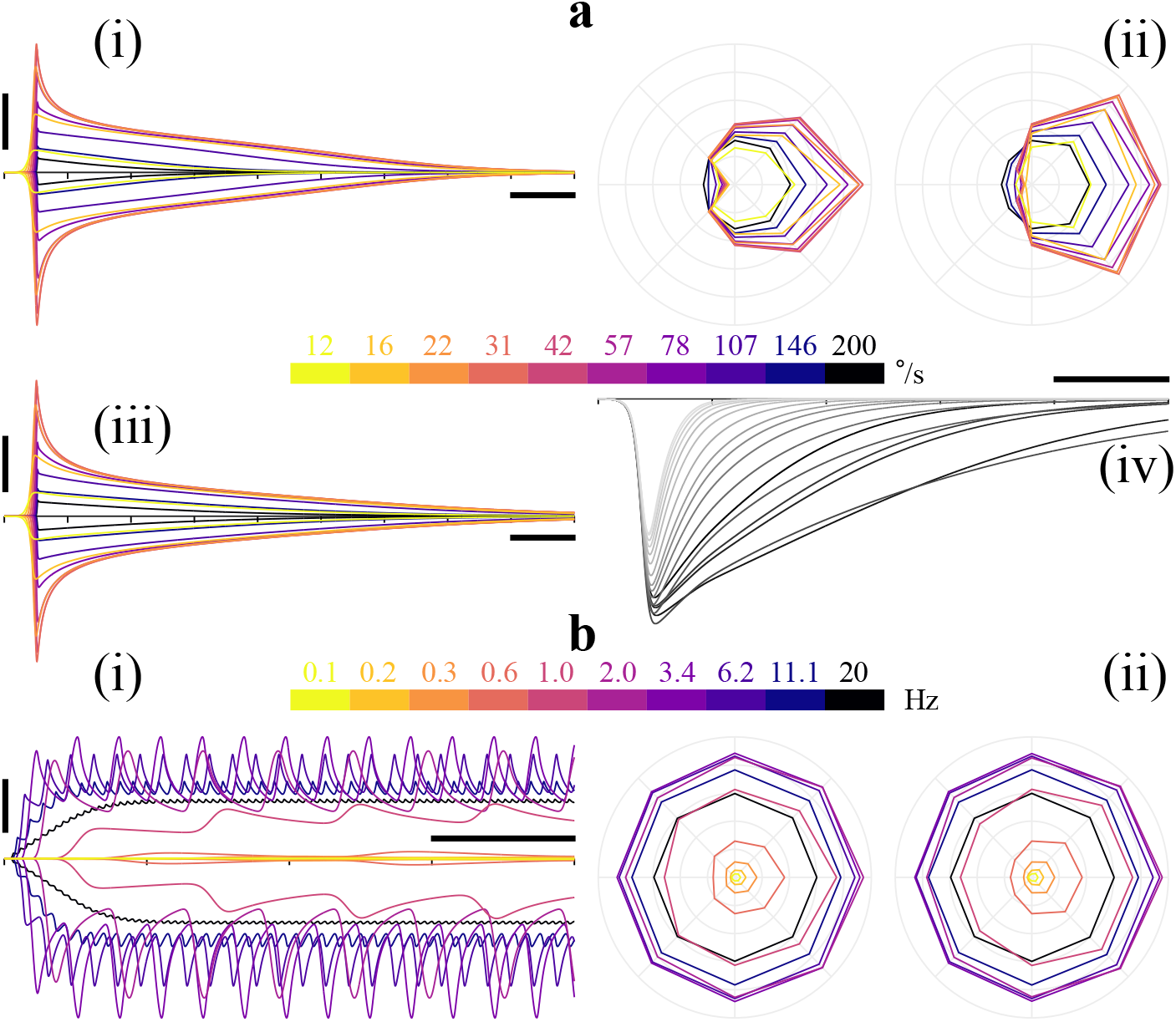
Responses of Mi4, Mi9, and T4 to bar and drifting sinusoidal grating stimuli. **a,** Bar stimuli. (i) Responses of Mi4 and Mi9 to a 2° wide bar traveling at different velocities. The responses are color coded and shifted along the ordinate by their respective equilibrium resting potentials. (ii) Corresponding responses of the home module T4s tuned to the two cardinal directions. Peak response above equilibrium resting potential is displayed for each stimulus direction PD, PD±*π/*4, PD±*π/*2, ND±*π/*4, and ND, for the different velocities. (iii) The Mi4-Mi9 dynamics is robust: response of a network with different synaptic parameters (Methods) from another optimization run. (iv) The optimization in action: the delayed and extended dynamics (only Mi9 shown) evolved gradually with changing synaptic parameters for the same stimulus. **b,** Sinusoidal grating stimuli. (i) Responses of Mi4 and Mi9 to a 20° wavelength sinusoidal grating traveling at different velocities. The responses are color coded and shifted along the ordinate by their respective pre-stimulus equilibrium resting potentials. Both the charging-up and the low-pass nature of the circuit are manifest. (ii) Same as a(ii) for this stimuli. Horizontal scale bars in a(i), a(iii), a(iv), b(i) = 1*s*. Vertical scale bar in a(i), a(iii) = 2*mV*, in b(i) = 3*mV*. Voltage increment of concentric circles in a(ii) = 2*mV*, in b(ii) = 1*mV*.

The resultant optimized behavior of Mi4 and Mi9 is, crucially, consistent with the neurons’ temporal kernels deduced from reverse correlation experiments [3]. Additionally, we found that although the temporally extended responses of the neurons permit simple synaptic integration of their signals at T4 to be sufficient to induce direction selectivity [13], both preferred direction enhancement and null direction suppression [14, 19] is necessary to abridge the overall response. Finally, the characteristics of the response to drifting sinusoidal gratings are readily explained by the charging-up of the recurrent networks and their low-pass nature (Fig.2b). We elaborate on these results below.

### The End-to-End EMD Circuit Implementation

Phototransduction in the Drosophila photoreceptor has been investigated extensively [21, 18]. We used the Wong and Knight model with its reported parameters [18] (order *n* = 8, *τ* = 2*ms*) without static nonlinearities to model the photoreceptor potential (Methods). The histaminergic photoreceptor-LMC synapse, responsible for the sign inverted response in L1 (Fig.1b-ii), has likewise been studied in detail [32], its high pass nature attributed to presynaptic subtraction of the variable extracellular field potential at the synapse [31]. We modeled the field potential as low pass filtered photoreceptor potential (*τ* = 2*ms*) and set the difference as the *V*_*pre*_ for the hyperpolarizing synapse (Methods).

The lengths, diameters, and synaptic polarities of L1, Mi1, Tm3, C3, L5, Mi4, Mi9, and T4 were drawn from [29, 22] (Table.1). L1 was assigned a 15*μm* high conductance initial synaptic zone [24]. Simulations were conducted using *l* = 30*μm* compartments with a 0.01*ms* time step size and potentials specified in *mV*. We verified that the results were identical to within precision bounds when using *l* = 10*μm* compartments (Fig.S1-2, S4). Notably, extensive simulations with randomly sampled diameters and lengths indicated that the neurons in the circuit were approximately equipotential across their entire cell body (Methods). The length of the neurites could not therefore be implicated in the generation of the requisite delays.

**Table 1:** Parameters for the full network using *l* = 30*μm* compartments.

| Presynaptic Neuron<br>(Diameter $d$, Length $l$) | Postsynaptic Neuron | $E_{syn}$ | $\mu$ | $\nu$ | $\eta$ |
| --- | --- | --- | --- | --- | --- |
| R1-R6 ($-, -$) | L1 | $-85mV$ | 40.0 | 3.0 | -3.82 |
| L1 ($1\mu m, 100\mu m$) | Mi1/C3/L5 | $-85mV$ | 5.0 | 0.4 | 27.4 |
| L5 ($0.6\mu m, 60\mu m$) | Mi4 | $0mV$ | 0.6 | 0.3 | 15.0 |
| C3 ($0.6\mu m, 60\mu m$) | Mi9 | $-85mV$ | 0.6 | 0.3 | 15.0 |
| Mi4 ($0.6\mu m, 60\mu m$) | Mi9 | $-85mV$ | 4.03555 | 0.29494 | 15.10545 |
| Mi9 ($0.6\mu m, 60\mu m$) | Mi4 | $-85mV$ | 4.03574 | 0.29485 | 15.10667 |
| Mi1 ($0.6\mu m, 60\mu m$) | T4a(b) | $0mV$ | 1.5 (1.5) | 0.1 | 7.4912 |
| Mi9 ($0.6\mu m, 60\mu m$) | T4a(b) | $-85mV$ | 1.5 (2.25) | 1.0 | 58.435 |
| Mi4 ($0.6\mu m, 60\mu m$) | T4a(b) | $-85mV$ | 1.5 (2.25) | 1.0 | 60.317 |

L1’s hyperpolarizing synapses onto Mi1, L5, and C3, responsible for their sign inverted back response, were modeled to be identical (Table.1) with *μ* set to fit the 10 − 20*mV* recorded response to flashes [5], and *η* set near saturation to induce the requisite half wave rectification (Fig.1b-ii and Methods). Since Mi1 and Tm3 have similar response profiles for moderate stimulus velocities, and both depolarize T4 [28], we chose to use only Mi1 in our parsimonious EMD model, consistent with the observed impact of silencing Tm3 [2, 28].

The mechanism that can potentially engender a delayed and extended response in a recurrent Mi4-Mi9 network can be conceptualized as follows. Mi4 is depolarized by L5’s synaptic input whereas Mi9 is hyperpolarized by C3’s synaptic input [29]. The reciprocal Mi4-Mi9 synapses are, however, both hyperpolarizing [28, 29]. A hyperpolarized Mi9 in turn anti-hyperpolarizes Mi4, while a depolarized Mi4 hyperpolarizes Mi9, triggering a positive feedback loop. With appropriately set *μ*’s, *ν*’s and *η*’s, this feedback can be tuned to both commence and return cells to equilibrium slowly. We searched for such parameters using the optimization procedure (Fig.2a-iv) that incrementally pushed the peak response time of Mi4 and Mi9 to the future, when the photoreceptors were driven by a bar stimulus. We found a wide range of parameters for which this was achievable (Fig.2a-i, iii and Fig.3a), demonstrating that the phenomenon was robust. Table.1 reports the parameters of the particular network presented here. Importantly, we found that the dynamics could be replicated using a single drive: eliminating either the L5 or C3 synapse and raising the *μ* of the other (Methods).

**Figure 3:**
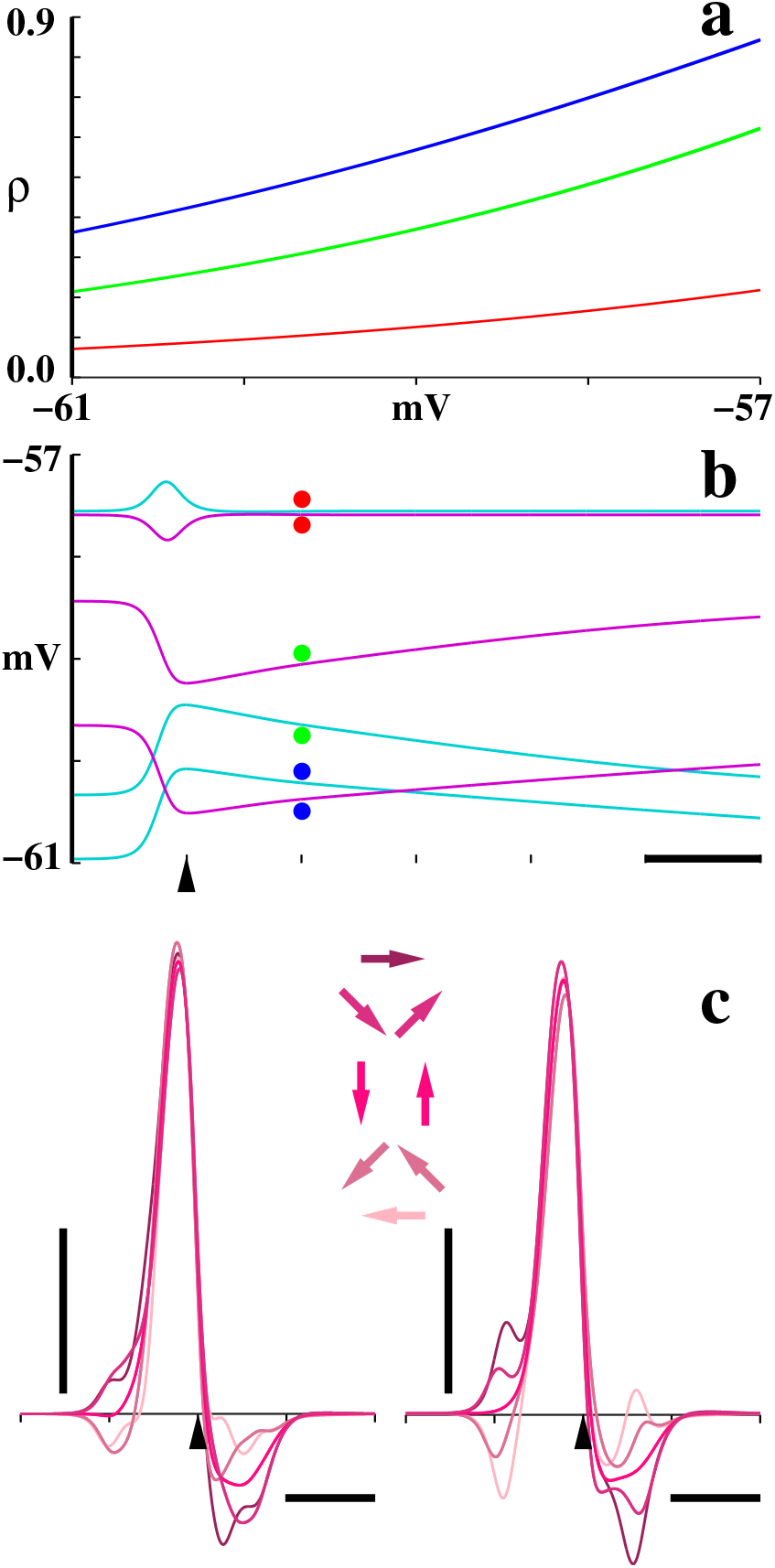
Pre- and Post Optimized Mi4-Mi9 network and corresponding T4 behavior. **a,** *ρ*(*V*_*pre*_) of the pair of synapses. Initialization, i.e., pre-optimized synapse pair labeled in red. Optimized synapse pair of the network presented in the article labeled in green. A second optimized pair, corresponding to the network in Fig.2a-iii labeled in blue. Note that the range [−61, −57]*mV* subsumes the operating range of all three synapse pairs as shown in (b). The optimization was not constrained to update the synapses symmetrically. However, it did to within precision bounds. All parallel graphs of *ρ*(*V*_*pre*_) between the green and blue curves manifest the delay necessary to implement the EMD, demonstrating that the phenomenon is robust. **b,** Responses of Mi4 (cyan) and Mi9 (magenta) in the three networks for a 2° wide bar of light traveling at 12°/*s*. Traces are labeled with adjacent color coded dots to distinguish between the three networks. Not only does the optimization delay the time to peak for both Mi4 and Mi9, it also substantially slows the tail decay. Note that both *V*_*rest*_ and *V*(*t*) change as the synaptic parameters are changed. **c,** Responses of the home module T4s tuned to the two cardinal directions (as in Fig.1b-i) when the Mi4/Mi9 synapses are in the pre-optimized parameter setting in red from (a). Compare these to Fig.1b-iii which corresponds to the Mi4/Mi9 synapses in the post-optimized parameter setting in green from (a). Stimulus directions PD, PD±*π/*4, PD±*π/*2, ND±*π/*4, and ND, are with respect to the insets in Fig.1b-i. Note that directionally selective response is almost absent. Horizontal scale bars in (b), (c), = 500*ms*. Vertical scale bar in (c) = 1*mV*. Pointers on abscissa mark time alignment for the home module as in Fig.1b.

We found that setting the peak response time of Mi4/Mi9 to be delayed by *≈* 125*ms* with reference to that of Mi1 (Fig.1b-ii) was sufficient to induce robust direction selectivity in T4 (Fig.1b-iii), owing to the slow decay (> 4*s*) of the Mi4/Mi9 response (Fig.2a-i). However, to abridge the T4 response, it was necessary to have the postsynaptic impact of the decays counteract one another symmetrically: we built a T4 cell that received a depolarizing synapse from the Mi1 of the home module and hyperpolarizing synapses from three Mi9s and three Mi4s [28] from the six neighboring modules (Fig.1a), such that a preferred direction (PD) stimulus triggered, in order, Mi9-Mi1-Mi4 and a null direction (ND) stimulus triggered them in reverse. The *μ*’s, *ν*’s, and *η*’s of the synapses were set to evoke commensurate effect from each synaptic input, operate in the linear regime, and be symmetric for each Mi4-Mi9 opposing pair to elicit cancellation (Table.1).

The robust directional response of T4 to bar stimuli traveling at > 100°/*s* [13] (Fig.2a-ii) seems at odds with its equilibrium response to drifting sinusoidal gratings. Grating associated direction selectivity of wide field horizontal cells, the postsynaptic target of T4, has been observed to be a function of temporal frequency [25]—grating velocity divided by wavelength— peaking at a mere 1*Hz*. We found that the Mi4-Mi9 network charges-up when driven by gratings, with respective baseline potentials equilibrating at values determined by the grating’s frequency (Fig.2b-i). Setting the sensitivity *ν* of the Mi4/Mi9 synapses on T4 to saturate/close at the peak of the 1*Hz* charged-up equilibrium response of Mi4/Mi9 (Methods) led the synapses to either be persistently open at near full conductance, or be closed, at > 1*Hz*. This resulted in the progressive eradication of grating associated directional responses at higher frequencies (Fig.2b-ii), without compromising bar stimuli associated responses (Fig.2a-ii).

## Discussion

### Synaptic Profiles can be the Key Determinant of Behavior

Recurrent connectivity is ubiquitous in animal brains. Without precise knowledge of the corresponding synaptic profiles, however, it is difficult to assess what role those recurrent connections play in the overall behavior of a given network. The optimized EMD circuit drives home this message definitively. Careful analysis of the dynamics of the network has shown us that the profiles of just two synapses, Mi4’s on Mi9 and Mi9’s on Mi4, in relation to the profiles of the other synapses, determine the emergence of the intended behavior of the network. Fig.3a presents the profiles *ρ*(*V*_*pre*_) of the pair of synapses over the operational range of the *V*_*pre*_ of Mi4 and Mi9, before and after optimization, and Fig.3b presents the corresponding voltage profiles of Mi4 and Mi9 when the photoreceptors are driven by the bar stimulus. As is clear from the voltage profiles of T4 in Fig.3c (as compared to that in Fig.1b-iii) directionally selective response is entirely absent in the pre-optimized network. The critical dynamical characteristics of a network can therefore be determined by the profiles of a select few constituent synapses; broad connectivity information is inadequate in such cases.

### Whence the Delay

Although the nonlinear dynamics of recurrent networks belie simplistic explanations in the general case, several fortuitous features of the optimized Mi4-Mi9 network allow for the identification of the source of its delayed response. First, the observation that the neurons are approximately equipotential across their entire cell bodies allows us to model both Mi4 and Mi9 as single compartments. Second, the profiles of *ρ*(*V*_*pre*_) for Mi4 and Mi9 in Fig.3a, for their respective operational ranges, suggest that they can be approximated by affine functions: *ρ*(*V*_*pre*_) = *a* + *bV*_*pre*_. Third, the symmetric dynamics of Mi4 and Mi9, as displayed in Fig.1b-ii and Fig.3b, permits the approximation 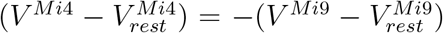. When the ODEs for Mi4 and Mi9 are simplified using these substitutions, one finds that the ODEs, shifted by respective equilibrium resting potentials, are identical to first approximation.

Equivalently, the Mi4-Mi9 recurrent network can be approximated as a single, one compartment, neuron with an *inverted autapse* following the now quadratic ODE:

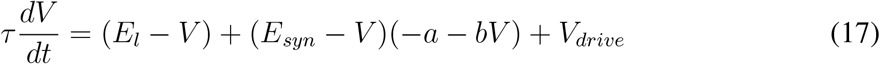

where *V*_*drive*_ approximates the L5 to Mi4 or C3 to Mi9 drive. Assuming equilibrium *V*_*rest*_ is at *V*_*drive*_ = 0, the right hand side of the reduced model Eq. 17 can the rewritten in terms of 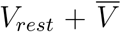 where 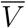 is the deviation of the potential from *V*_*rest*_. We then get 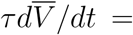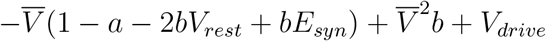. To a first approximation, the time constant near equilibrium therefore scales by 1/(1 − *a* − 2*bV*_*rest*_ + *bE*_*syn*_), which depending on the values of *a, b, V*_*rest*_, *E*_*syn*_ can be very large, causing the response to have a slow rise and slow decay.

### A More Nuanced Description of the EMD Mechanism

The HR and BL models prescribe delay lines that are nonlinearly integrated with the direct lines to engender direction selectivity. We have however found that the mechanism that leads to direction selectivity in the model EMD neural circuit is more subtle and emerges by way of several interacting processes. The delay originating from the nonlinear dynamics of the recurrent Mi4-Mi9 network comes with an associated broadening of the Mi4/Mi9 response (Fig.1b-ii). The magnitude and width of this response (Fig.2a-i and iii) vary nonlinearly with the velocity of the input stimulus. The benefit of this broadening of response is that it functions as an additional delay. This is why a delay of *≈* 125*ms* between the peak response time of Mi1 and that of Mi4/Mi9 was found to be sufficient to induce robust direction selectivity, even when the peak to peak delay between the profiles of neighboring photoreceptors in the lattice ranged between 212 to 368*ms* (Fig.1b-i). The drawback of this broadening is that the tail of the response can persist for longer than is desirable. The EMD circuit eliminates the tail response via opponency: the Mi4 and Mi9 of diametrically opposed lattice points feed the home network. The result is that for a PD stimulus the Mi1 response of the home module sits atop a depolarizing bump generated by the Mi9-then-Mi4 response. Contrastingly, for a ND stimulus the Mi1 response sits atop a hyperpolarizing dip generated by the Mi4-then-Mi9 response. Simple synaptic integration of these signals at T4 therefore suffices to induce direction selectivity.

Our model has thus shown that recurrent dynamics can explain the EMD. Importantly, the optimization procedure that we have leveraged to arrive at the model parameters holds promise for networks beyond the one addressed here.

## Materials and Methods

Custom code written in C was used both for simulating the dynamics of the model as well as for the optimization. The programs were run on a 10-Core, 2.3GHz Linux server with 64 GB RAM.

### Compartment Model

Each compartment *i* was modeled to be cylindrical of diameter *d*^*i*^ and length *l*^*i*^, following 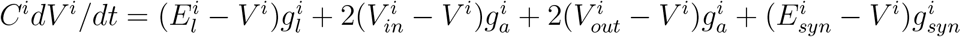, (Fig.1c) with the differential equation divided on both sided by 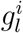 to ease the derivation of parameters as well as to ensure numerical stability. Universal membrane constants [12] were set as: specific membrane capacitance *C*_*m*_ = 1*μF/cm*^2^, specific membrane resistance *R*_*m*_ = 20*K*Ω*cm*^2^, and specific axial resistance *R*_*a*_ = 100Ω*cm*. For a cylindrical compartment *i* with surface area *A*^*i*^ and cross sectional area *a*^*i*^, *C*^*i*^ = *C_m_ × A^i^*, 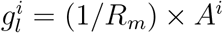, and 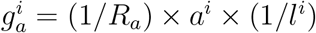. Parameters were first computed for a canonical cylinder *i* = 0 of diameter *d*^0^ = 1*μm* and length *l*^0^ = 100*μm*. For all compartments *i*, 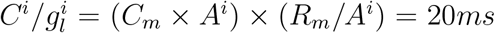. For canonical cylinder *i* = 0, calculations yield 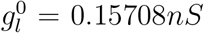, and the unitless 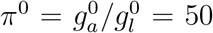. Hence for cylinder *i*, 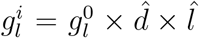, and 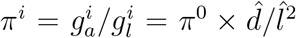, where 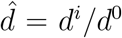 and 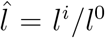. The unitless time varying 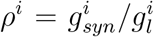 which is a function of *V*_*pre*_—the *V*^*i*^ of the presynaptic compartment—was modeled as 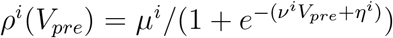. *ρ*^*i*^ therefore ranged between 0 and *μ*^*i*^ depending on *V*_*pre*_. *E*_*l*_ was set at −55*mV*, and *E*_*syn*_ at −85*mV* for hyperpolarizing and 0*mV* for depolarizing synapses.

The entire differential-algebraic system of equations was simulated using the forward Euler method. Simulations were initially conducted with compartments of length *l* = 10*μm* and a time step size of 0.001*ms*. However, it was observed that for the range of diameters used, the results were identical to within precision bounds with *l* = 30*μm* compartments simulated at a 0.01 *ms* time step size (Fig.S1 and S2).

Two idiosyncrasies of tonically active synapses were immediately evident. Firstly, the equilibrium resting potential of a neuron varied markedly [12] depending on how conductive its input synapses were at equilibrium, which then depended on the *μ*, *ν*, *η* and equilibrium resting potential of each of its presynaptic neurons. Secondly, appropriate adjustments to *μ*, *ν* and *η* could replicate the qualitative dynamics for different settings of *E*_*l*_ and *E*_*syn*_.

### Morphology of individual Neuron Models

The Drosophila L1 has diameter *d* ≈ 1*μm* [22] and length *l* ≈ 100*μm* with an initial high conductance synaptic zone. We modeled a 15*μm* synaptic zone [22] for which *g*_*l*_ and the equilibrium *g*_*syn*_ were scaled up ten-fold [24]. A *d* = 1*μm*, *l* = 15*μm*, high conductance synaptic zone compartment for L1 was modeled as follows. Since *g*_*l*_ scales by (*dl*) and *g*_*a*_ scales by (*d*^2^*/l*), to have *g*_*l*_ increase ten-fold [24] while maintaining the value of *g*_*a*_ fixed, *d* for the compartment was scaled ×10^1/3^ and *l* was scaled ×10^2/3^, to the new values *d* = 2.1544347*μm* and *l* = 69.623832*μm*.

When this L1 synaptic compartment was linked to a 100*μm* axon built out of *d* = 1*μm*, *l* = 10*μm* compartments, we found little difference between the *V*^*i*^ at the synaptic versus the most distal compartment (Fig.S1). We conducted a large set of simulation experiments where the diameters of the ten compartments were chosen randomly from an uniform distribution over the range [−0.7, 1.3]*μm*. In all cases, we found little difference between the *V*^*i*^ at the synaptic versus the most distal compartment (almost identical to Fig.S1 and hence not shown). Nor did we find a difference when appropriately fewer compartments of longer *l* were used (Fig.S2). Even with a 500*μm* axon built out of *l* = 10*μm* compartments, the distal *V*^*i*^ was delayed by a mere 5*ms*, although the equilibrium *V*^*i*^ value shifted closer to *E*_*l*_ (Fig.S3). In the final model, L1 was built out of three compartments linked in series: the high conductance synaptic zone compartment, an intermediate *d* = 1*μm*, *l* = 60*μm* compartment, and a distal *d* = 1*μm*, *l* = 40*μm* compartment. The *V*^*i*^ of the distal compartment was set as the *V*_*pre*_ of L1’s synapses on Mi1, C3, and L5.

The Drosophila C3 has diameter *d* ≈ 0.63*μm* [22]. For C3, Mi1, L5, Mi4, and Mi9, we set each *d* = 0.6*μm*, and since they span the ~ 50*μm* medulla [29], we set each *l* = 60*μm*. The cells Mi1, C3, L5, Mi4, and Mi9 were all built to be identical in their morphology: each neuron was built as two compartments each of *d* = 0.6*μm*, *l* = 30*μm*, linked in series. We verified that the recurrent Mi4-Mi9 network had identical properties when they were each assembled out of six *l* = 10*μm* compartments (Fig.S4). The same was true for Mi1, C3 and L5. Finally, the entire network was replicated (Fig.1a) for each of the seven contiguous modules in the hexagonal lattice.

T4 was built out of seven compartments each of *d* = 0.6*μm*, *l* = 30*μm*. The home compartment was linked to each of three compartments at one terminal and each of three other compartments at the other. The home compartment received a synaptic input from the Mi1 of the home module; each of the remaining six compartments received a synaptic input from either the Mi4 or the Mi9 from a distinct surrounding module in the hexagonal lattice (Fig.1a).

### Photoreceptor and the Photoreceptor-L1 synapse

The Drosophila has neural superposition eyes where each module in the lamina receives the axon of one of each photoreceptor R1-R6, all of which manifest the same visual axis [1]. Since each omatidium contains one of each R1-R6, the angle between the visual axes of the R1-R6 in an omatidium matches the angle between the axes of neighboring omatidia [7]. Hence the visual axis of neighboring modules in the lamina are separated by the same angle as that of the omatidia. The omatidia in the Drosophila eye are arranged with interomatidial angle of ≈ 5.1° [27]. The normalized angular sensitivity of R1-R6 in each omatidium is well approximated by 2-D isotropic Gaussians with full width at half maximum (FWHM) estimated at 4.2° when the combined effects of geometric and wave optics are invoked [27]. Hence a photoreceptor centered at 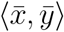 has normalized angular sensitivity 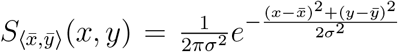, where 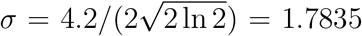. We modeled seven contiguous omatidia as individual photoreceptors centered on lattice points 5.1° apart on a hexagonal lattice (Fig.1b-i), with angular sensitivity 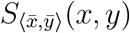. For any given time varying input stimulus *I*(*x, y, t*), the input drive for the photoreceptor centered at 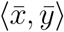 was computed as 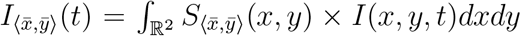. For both the bar and the drifting sinusoidal grating stimuli, the X-Y plane of integration was first rotated so as to align one of the axes to be orthogonal to the stimulus direction. This allowed the integral to be simplified in closed form to a 1-D integral over ℝ. Numerical integration was then performed with a spatial step size of 0.04°.

The Drosophila photoreceptor response has previously been modeled as a NLN (nonlinear static-linear dynamic-nonlinear static) cascade, where the two static nonlinearities are third order polynomial functions and the linear component is the Wong and Knight photoreceptor model 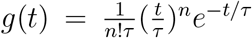, with the model parameters fit to recorded data [18]. Since the two polynomial fits were found to be fairly close to linear for wild type Drosophila [18], we used only the linear component with the parameters [18] *n* = 8 and *τ* = 2*ms*. For a photoreceptor centered at 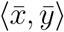, the photoreceptor potential was therefore modeled as 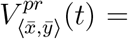 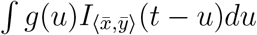.

The photoreceptor-LMC synapses being histaminergic, we set *E*_*syn*_ = −85*mV* for the photoreceptor-L1 synapse. It has been demonstrated [31] that at this specialized synapse, the extracellular field potential changes with illumination, and the presynaptic transmembrane potential in the photoreceptor axon is well approximated by subtracting the field potential from the intracellular potential. We modeled the field potential as low-pass filtered photoreceptor potential: *τdV^fp^/dt* = −*V^fp^* + *V^pr^* with *τ* = 2*ms*, and set the *V*_*pre*_ of the photoreceptor-L1 synapse as *V*_*pre*_ = *V ^pr^* − *V^fp^*.

### Synaptic Parameters

Key to most of the parameter settings are the observations that 1/(1 + *e*^−*x*^) is convex for *x* < 0, concave for *x* > 0, approximately linear over *x* ∈ [−1.5, 1.5], and saturates beyond *x* = ±5. For all feedforward synapses, a range of *x* was first chosen that induced the desired output of *ρ*, following which the scale and shift parameters *ν* and *η* were computed so as to match it to *V*_*pre*_’s range. The following parameters of the photoreceptor-L1 synapse relate to a stimulus intensity for which a 2° wide bar traveling at 12°/s generated a *V*_*pre*_ = *V^pr^* − *V^fp^* at the synapse of range ±0.33*mV* about a baseline of 0*mV*. Since the range of the *V*_*pre*_ scales linearly with the intensity, the value of *ν* can be scaled to evoke the same postsynaptic response for a different intensity value. We set ⟨*μ*, *ν*, *η*⟩ = ⟨40, 3, −3.82⟩ for the photoreceptor-L1 synapse. The resultant postsynaptic response to the bar in the high conductance synaptic compartment was a hyperpolarization of 6.98*mV* followed by a depolarization of 5.31*mV* about an equilibrium resting potential of −65.22*mV* (Fig.1b-ii). The *η* offset was chosen to model the slightly larger hyperpolarization reported in experiments [32]. As noted earlier, the entire three-compartment L1 was equipotential to within 0.3*mV*.

L1’s synapse on Mi1, C3, and L5 were modeled to be identical. Since the synapses are hyperpolarizing, we set *E*_*syn*_ = −85*mV*. We set ⟨*μ*, *ν*, *η*⟩ = ⟨5, 0.4, 27.4⟩, which for the noted bar stimulus resulted in a depolarization of 10*mV* followed by a much smaller hyperpolarization of 1.3*mV* about an equilibrium resting potential of −74.97*mV* (Fig.1b-ii). The values of *μ* and *ν* were set to mimic the response of Mi1 to flash stimuli [5]. The value of *η*, set close to saturation at the equilibrium *V*_*pre*_ = −65.11*mV* of L1’s distal compartment, yielded the intended half wave rectification.

L5’s synapse on Mi4 being depolarizing and C3’s synapse on Mi9 being hyperpolarizing, we set their *E*_*syn*_ = 0*mV* and −85*mV*, respectively. Both synapses were assigned ⟨*μ*, *ν*, *η*⟩ = ⟨0.6, 0.3, 15⟩. We found that either synapse could be eliminated and the other set at ⟨*μ*, *ν*, *η*⟩ = ⟨2, 0.3, 15⟩ to replicate the dynamics in Fig.2a-i. The reciprocal synapses between Mi4 and Mi9 are both hyperpolarizing, hence we set both *E*_*syn*_ = −85*mV*. Their synaptic parameters were then arrived at via the optimization procedure. We found a large range of parameter settings for which the Mi4-Mi9 recurrent dynamics was delayed and extended (Fig.2a-i and iii). The model reported here (Fig.2a-i, Table.1) has ⟨*μ*, *ν*, *η*⟩ = ⟨4.03555, 0.29494, 15.10545⟩ for Mi4’s synapse on Mi9, and ⟨*μ*, *ν*, *η*⟩ = ⟨4.03574, 0.29485, 15.10667⟩ for Mi9’s synapse on Mi4. For the noted bar stimulus, this resulted in Mi4 depolarizing by 0.88*mV* about an equilibrium resting potential of −60.32*mV*, and Mi9 hyperpolarizing by 0.81*mV* about −58.44*mV* (Fig.1b-ii). The delayed and extended dynamics evolved gradually with changing parameters during the optimization procedure (Fig.2a-iv). The stated parameter values were chosen to have the peak response of Mi4/Mi9 be delayed by *≈* 125*ms* with reference to that of Mi1. A different optimization run that achieved an even larger delay (Fig.2a-iii) gave ⟨*μ*, *ν*, *η*⟩ = ⟨2.34727, 0.28076, 15.42189⟩ for Mi4’s synapse on Mi9, and ⟨*μ*, *ν*, *η*⟩ = ⟨2.35028, 0.28072, 15.42252⟩ for Mi9’s synapse on Mi4.

Mi1’s synapse on T4 being depolarizing, we set its *E*_*syn*_ = 0*mV*. Mi4’s and Mi9’s synapses on T4 being hyperpolarizing, we set their *E*_*syn*_ = −85*mV*. The home module Mi1’s synapse on T4 was set at ⟨*μ*, *ν*, *η*⟩ = ⟨1.5, 0.1, 7.49⟩. For cardinal direction ‘a’ (Fig.1b-i), the three Mi9 synapses on T4—from Mi9s of three contiguous modules—were set at ⟨*μ*, *ν*, *η*⟩ = ⟨1.5, 1.0, 58.44⟩. Additionally, the three Mi4 synapses on T4—from Mi4s of the three symmetrically opposing contiguous modules—were set at ⟨*μ*, *ν*, *η*⟩ = ⟨1.5, 1.0, 60.32⟩. For cardinal direction ‘b’ (Fig.1b-i), the *μ*’s were raised to 2.25 along the PD-ND axis and dropped to 0 orthogonal to it. The *μ*’s in both cases were set to mimic the reported response of T4 to the noted bar stimulus [13]. The *ν*’s for the Mi4 and Mi9 synapses on T4 were set such that at their charged-up peak response to a 1*Hz* drifting sinusoidal grating, amounting to a shift of *≈* 2.8*mV* (Fig.2b-i), the synapses reached saturation and closure, respectively.

### Equilibrium Resting Potentials

We began by setting the photoreceptor potential to a constant 0 over all time. The network is at its equilibrium resting state when *dV*^*i*^*/dt* = 0 for all compartments *i*. Setting *dV*^*i*^*/dt* = 0 for all *i* turns the differential-algebraic system of equations into an algebraic system of equations. We initialized this system at 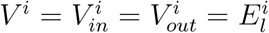 for all *i*, and solved it using the fixed point Jacobi iteration scheme. The iterations were continued until convergence, set as all new values evaluating to within 10^−9^*mV* of their previous values.

---

**Supplementary Figure 1:**
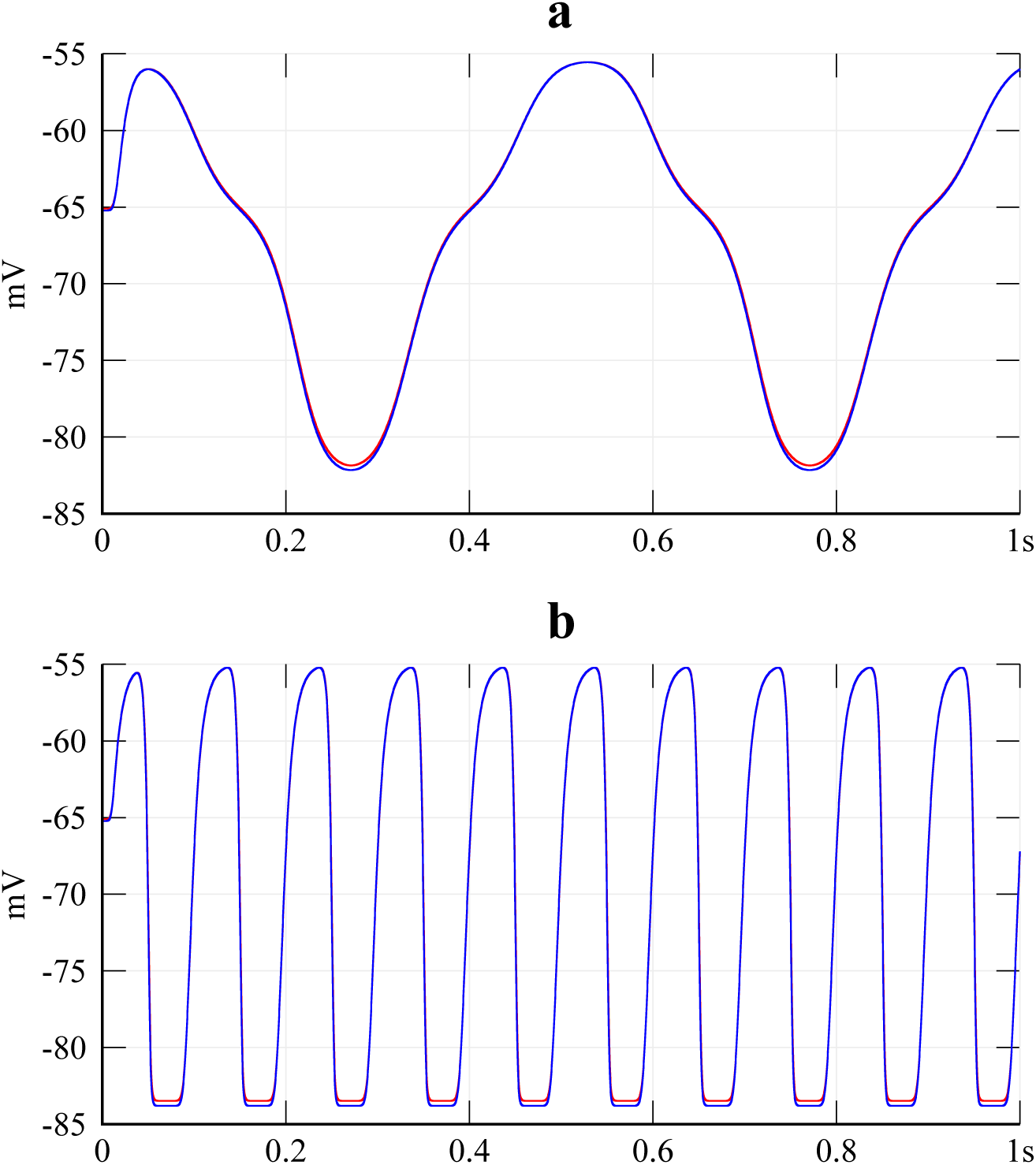
Neurons were almost equipotential across their entire cell body. A *d* = 1*μm*, *l* = 115*μm* L1, built from ten *l* = 10*μm* and one *l* = 15*μm* high conductance synaptic zone compartment. Responses of the most distal of the ten (red) and the initial synaptic compartment (blue), to a wavelength = 20° square wave traveling at **a**, 2°/*s* and **b,**10°/*s*. Responses of all other compartments were in between. All other neurons, modeled with *d* = 0.6*μm*, *l* = 60*μm*, were even closer to being equipotential due to their larger axial to leak conductance ratios.

**Supplementary Figure 2:**
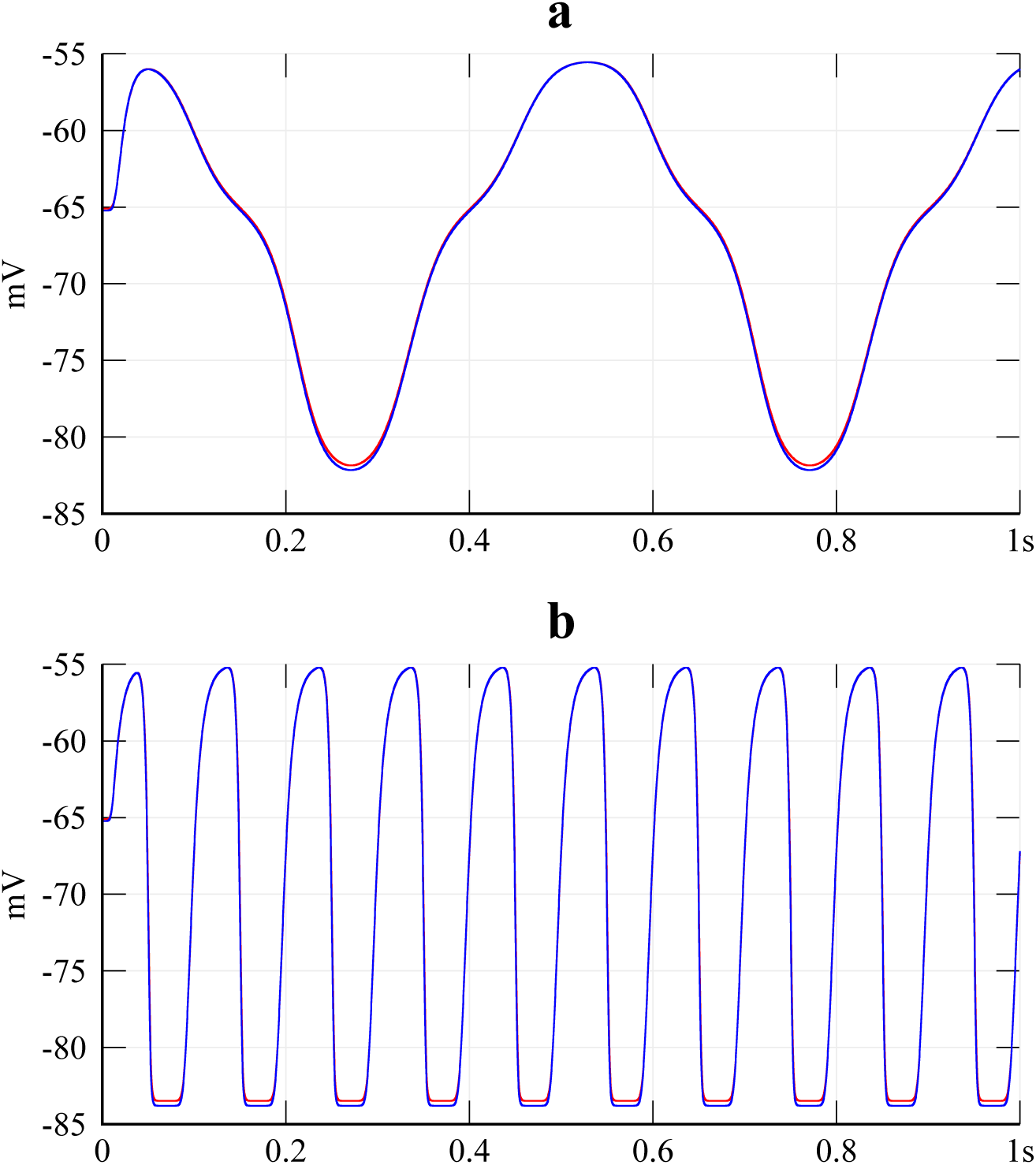
Assembly using fewer longer compartments is viable. The L1 in Fig.S1 built from a *l* = 40*μm* linked to a *l* = 60*μm* linked to the *l* = 15*μm* high conductance synaptic zone compartment. Responses of the *l* = 40*μm* (red) and the initial synaptic compartment (blue), to the same square wave stimuli in Fig.S1, i.e., traveling at **a**, 2°/*s* and **b,** 10°/*s*. To aid in comparison, we note that the responses of the high conductance synaptic zone compartment (blue) are identical to those in Fig.S1.

**Supplementary Figure 3:**
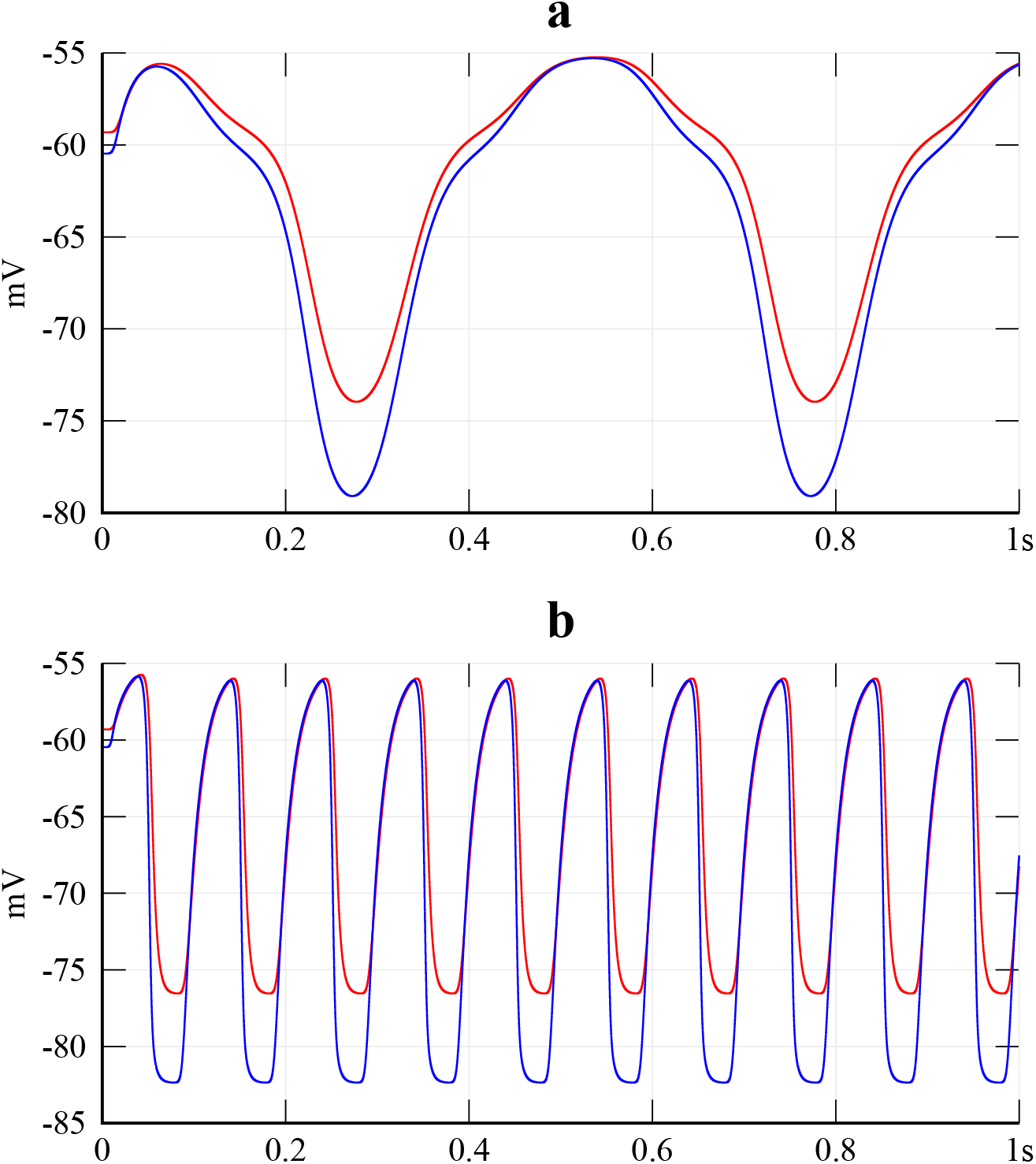
Cell body does not generate the requisite delay. A *d* = 1*μm*, *l* = 515*μm* L1, built from fifty *l* = 10*μm* and one *l* = 15*μm* high conductance synaptic zone compartment. Responses of the most distal of the fifty (red) and the initial synaptic compartment (blue), to a wavelength = 20° square wave traveling at **a**, 2°/*s* and **b,** 10°/*s*. Responses of all other compartments were in between. Although the responses are shifted toward *E*_*l*_ = −55*mV*, the peak to peak delay between the proximal and distal compartments is *≈* 5*ms*.

**Supplementary Figure 4:**
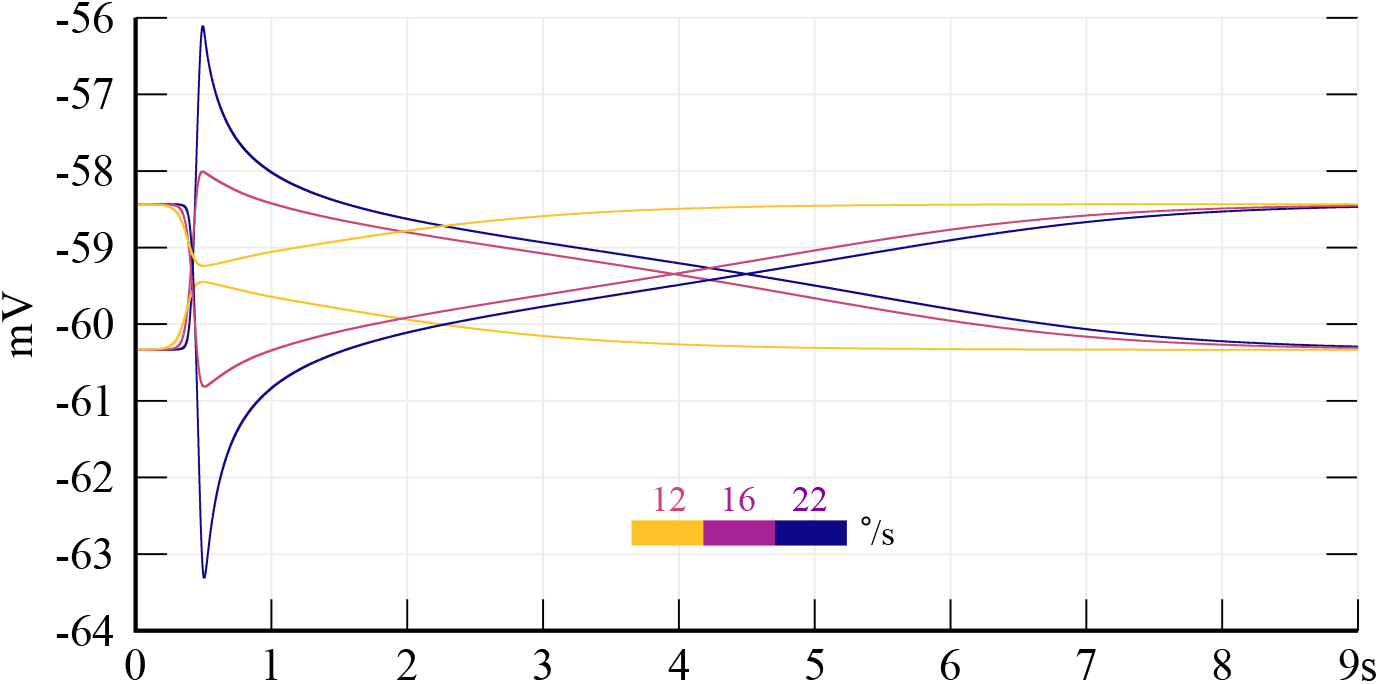
Mi4-Mi9 network response was identical to when neurons were assembled using shorter compartments. Neurons C3, L5, Mi4, and Mi9, modeled as *d* = 0.6*μm*, *l* = 60*μm*, were each built from six *l* = 10*μm* compartments. The responses were identical to within precision bounds as compared to the models where the neurons were each built out of two *l* = 30*μm* compartments. Raw responses of Mi4 and Mi9 are shown for a subset of velocities of the bar stimuli, for comparison against Fig.2a-i.

## Data availability

The data that support the findings of this study are publicly and openly available at https://doi.org/10.6084/m9.figshare.c.4503218.v2

